# Fine-Grained Detection of AI-Generated Writing in the Biomedical Literature

**DOI:** 10.64898/2026.01.01.697311

**Authors:** Richard She

**Affiliations:** School of Biological Sciences, Nanyang Technological University, Singapore

## Abstract

Generative AI systems are rapidly being integrated into scientific workflows, yet the specific ways in which AI-generated prose appears in published literature remain poorly characterized. Here, we use Pangram, a transformer-based detector optimized for adversarial paraphrasing, to analyze full-length biomedical research articles from 13 major journals. Papers published in 2021-2024 showed almost no detectable AI-generated text, whereas manuscripts published in 2025 exhibited a sharp increase, with 12.4% containing at least one localized passage classified as AI-written. AI usage was highly nonuniform across authors and geography: 32% of papers originating from South Korean institutions and 26% papers from Chinese institutions contained AI-generated passages, compared to 7.4% from U.S. institutions. In a focused case analysis, six labs that published fully AI-generated manuscripts also produced additional papers with extensive AI-generated segments. Journals likewise differed, with high-selectivity venues rarely containing AI-authored prose, while high-volume journals accounted for most AI-positive manuscripts. Together, these findings provide the first detailed empirical map of how and where AI-generated writing is entering the scientific literature, underscoring the need for clear norms and policies governing the use of generative AI in scientific communication.

## Introduction

*“It takes all the running you can do, to keep in the same place*.*”* – The Red Queen

At this point, everyone knows that their students are using AI to write their essays and do their homework. Who among us has not received an email beginning with “Dear Professor [Last Name].” For astute observers of more skilled students, one can still pick up on telltale AI signatures, from the proliferation of punctuation such as the em dash to an enrichment for uncommon verbs like “delve.” Furthermore, modern large language models (LLMs) are irresistibly drawn to a certain rhetorical tic, the corrective pivot, using an “not X but Y” cadence, no matter how you prompt them. It’s not a glitch of style — it’s actually their native register.

However, for any individual student, it is effectively impossible to prove guilt beyond a reasonable doubt. Because of its amorphous nature, AI-generated text has proven exceedingly difficult to watermark. Research at OpenAI on statistical watermarking stretches back to 2022^1^, but these schemes do not survive adversarial editing, and any forensic signal evaporates as soon as the text is retouched. In this landscape, the arms race is asymmetric, and the students are winning. With a bit of paraphrasing and cosmetic tweaking, they can assume impunity. Furthermore, the students are not alone; an informal poll of my former colleagues revealed that nearly every grad student and postdoc uses frontier LLMs every day. While LLM adoption likely varies by geography, age, and scientific field, given both the seductive power and genuine utility of these systems, one might reasonably wonder to what extent AI-generated writing has already made its way into the scientific literature.

Since individual cases of AI usage have thus far proven impossible to adjudicate, existing studies have focused on coarse grained metrics. One study that analyzed over 15 million abstracts from the biomedical literature documented the emergence of a bona fide LLM lexicon, with specific model-favored words rising at rates that cannot be explained by natural linguistic drift^2^. Other case reports point to the use of generative AI during peer review^3^, suggesting that academics find these tools both useful and embarrassing, deploying them only behind the shield of anonymity for work that never becomes part of the permanent record. A broader analysis of conference peer-review data reveals that the estimated fraction of LLM-generated reviews spikes as deadlines approach^4^, underscoring the messy practical and behavioral variables that determine where and when generative AI is used. However, these studies leave several critical questions unanswered: 1) who exactly is using generative AI? 2) to what extent is AI being used in fully published and peer-reviewed manuscripts? and 3) as AI assisted writing becomes more prevalent and inevitable, how will the scientific community choose to govern its use?

To answer these questions, we need tools that can surpass the limits of human intuition alone in detecting AI generated text^5,6^; in practice, it takes a neural network to reliably catch another neural network. Furthermore, state-of-the-art AI detectors must be robust against not only light paraphrasing but the next stage of the arms race: AI “humanizers”, a class of software tools designed to rewrite model output so that it appears plausibly human. Such humanizers break most naïve AI detectors^7^. We therefore turn to Pangram, a transformer-based neural network explicitly engineered for this adversarial setting. In head-to-head benchmarks it reaches approximately 99–99.8% accuracy with false-positive rates on the order of 0.01–0.1% across domains, including in scientific writing and essays by non-native English speakers, and remains highly effective even against adversarial “humanizer” paraphrases that are explicitly optimized to evade detection^8^. Crucially, Pangram operates on overlapping windows of text rather than whole documents, allowing it to flag localized AI-written segments embedded within otherwise human-authored manuscripts. In small scale testing, a recent Nature news report on Pangram’s deployment at the American Association for Cancer Research notes that a sizeable minority of peer-review reports and a smaller but non-trivial fraction of manuscripts contain detectable LLM text, almost always without disclosure^9^. Here we use Pangram to screen a large selection of biomedical research papers, both validating its performance in this domain and characterizing how, and where, AI writing is already seeping into the scientific literature.

## Results

The release of ChatGPT in late November 2022 provides a natural experiment for assessing Pangram’s AI-detection performance on biomedical manuscripts published before and after the widespread availability of large language models. To survey a broad cross-section of the literature, we first collected full-length articles from 13 well-known biomedical journals. As a negative control, we randomly sampled 1,000 papers published in 2021 and evaluated each for AI-generated text, excluding materials and methods sections and other ancillary segments outside of the main text. Reassuringly, all 1,000 manuscripts were classified as fully human-written (Figure 1A).

**Figure 1:**
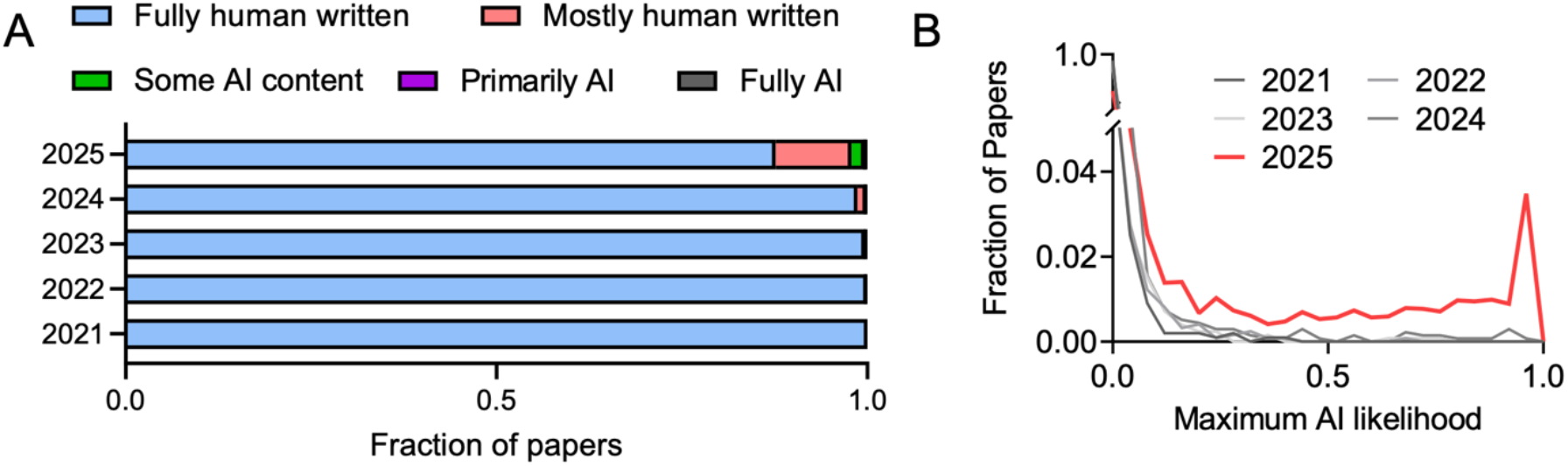
(A) Bar graph of the fraction of papers with AI-detected writing from 2021 to 2025. (B) Histogram of the maximum AI likelihood score within each paper for papers from 2021-2025.

As a positive control, we randomly selected 10 fully human-written papers from 2021 and removed either the abstract, introduction, or discussion sections. We then asked AI models to rewrite the lost section, while attaching the rest of the paper as a part of the prompt. For ChatGPT5 Thinking Mode, Gemini 3 Pro and Claude Opus 4.5, all resulting abstracts, introductions, and discussion sections were classified as fully AI-generated. However, for OpenAI’s latest update to ChatGPT5.1, released on November 12^th^ and thus not fully integrated into Pangram’s adversarial detection model, only 7 out of 10 abstracts were flagged as fully AI-generated. For the longer sections, 8 out 10 introductions and 9 out of 10 discussions were flagged as fully AI-generated. Within the 2 remaining introductions and 1 remaining discussion, Pangram’s sliding window analysis, which computes a per-passage AI likelihood in chunks of several hundred words, flagged the majority of windows as AI generated, with only a single plausibly human section with AI likelihood ranging from 0.25 to 0.43. By contrast, our entire dataset of 2021 human-written papers contained zero windows exceeding a 50% AI probability, far below Pangram’s standard detection threshold. These results highlight the remarkable false positive and false negative rates of Pangram’s detection tool, while also demonstrating its robustness to extremely recent model updates such as Gemini 3 and Opus 4.5. The only detectable slippage arises when a new frontier model introduces a genuinely novel stylistic regime, highlighting that as these systems evolve, any detector, no matter how capable, will be forced to chase multiple moving targets.

When we repeated this analysis for papers published from 2022–2024, we found only vanishingly rare instances of AI-generated content. However, an expanded sample of manuscripts published in 2025 showed a dramatic increase in AI usage (Figure 1A), with up to 12.4% of papers containing at least one window flagged as AI-generated. In most such cases, AI usage was confined to a small number of localized sections embedded within an otherwise human-written manuscript, though we also identified several papers classified as predominantly or even fully AI-generated. In aggregate, 2025 papers also exhibited a higher frequency of intermediate AI likelihood scores (Figure 1B), with the most straightforward interpretation of this trend being that 2025 manuscripts more commonly incorporated AI-generated text at initial stages but subsequently revised the text through multiple rounds of human editing and rephrasing. Taken together, these results indicate that the published biomedical literature is a lagging indicator of underlying author behavior, given typical publication delays of a year or more, and that the field now appears to be on the cusp of a substantial wave of AI adoption.

To quantify the geographic distribution of papers with AI generated writing, we mapped the institutional affiliations of each senior author in our 2025 dataset. This analysis revealed a significant enrichment for AI usage in papers originating from countries where English is not the primary native language. Among these, institutions in South Korea and China had the highest proportions of papers containing AI generated content, at 32% and 26% respectively (Figure 2). In contrast, only 7.4% of papers originating from institutions in the United States exhibited signs of generative AI. Among the U.S.-affiliated manuscripts, 20 contained AI-positive text in more than a single window, suggesting substantial AI involvement. Manual inspection of first author names suggested that all 20 of these papers had first authors with non-Anglophone names; 10 had Chinese given and family names without the anglicization that is common in American-born Chinese. While such name-based inference is necessarily imperfect, this pattern is consistent with generative AI being used preferentially by non-native English speakers as a key tool for scientific writing. In this sense, large language models may partially alleviate long-standing competitive disadvantages faced by scientists writing in a second language, even as they raise new questions about transparency, authorship, fairness, and the integrity of the scientific record.

**Figure 2:**
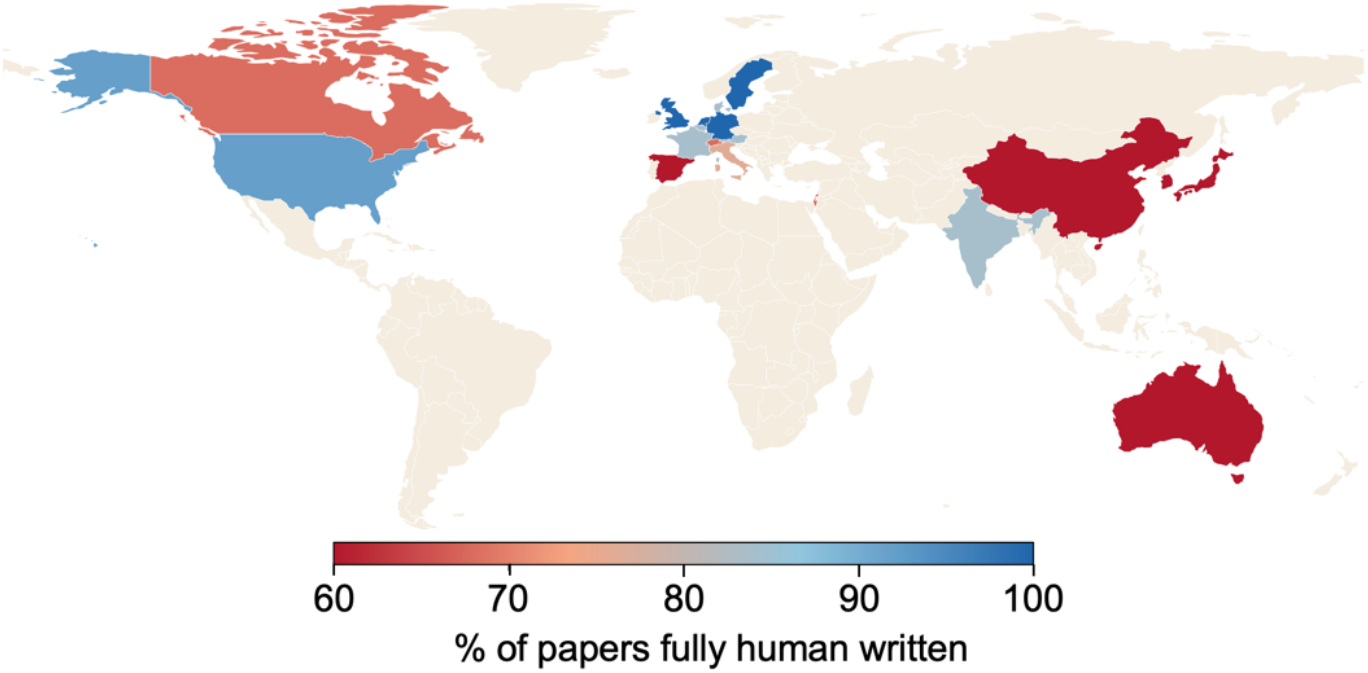
Geographic distribution of papers with AI generated writing.

In the most extreme cases, we detected 6 papers out of 5,077 that appeared to be fully AI generated. We manually confirmed that none of these manuscripts disclosed any use of generative AI in their Methods or Acknowledgements sections, despite explicit journal policies mandating such disclosure. To determine whether this reflected an isolated oversight by a busy PI or a consistent pattern of behavior, we contact traced all 2025 manuscripts published by the senior authors of these six papers. Among these 18 new papers, which were not included in our original analysis, only 2 were classified as fully human written (Figure 3). The remainder included 1 additional fully AI generated paper, 3 additional predominantly AI-generated papers, 8 papers with multiple AI-positive windows, and only 4 papers that could be classified as mostly human with some AI content detected. Although prior work has suggested that some AI detectors may falsely flag non-native English writing as AI-generated^10^, our manual inspection of these manuscripts indicates that this is unlikely to account for the systemic patterns observed. Instead, these results reveal a clear clustering of generative AI usage at the level of individual labs: most senior authors appear to avoid AI-generated text entirely, whereas others show consistent and repeated incorporation of AI-generated passages across multiple publications. This lab-specific bimodality provides a strong internal validation of Pangram’s detection consistency, and suggests that early adoption of generative AI in scientific writing is emerging not as random background noise but as distinct behavioral regimes within the research community.

**Figure 3:**
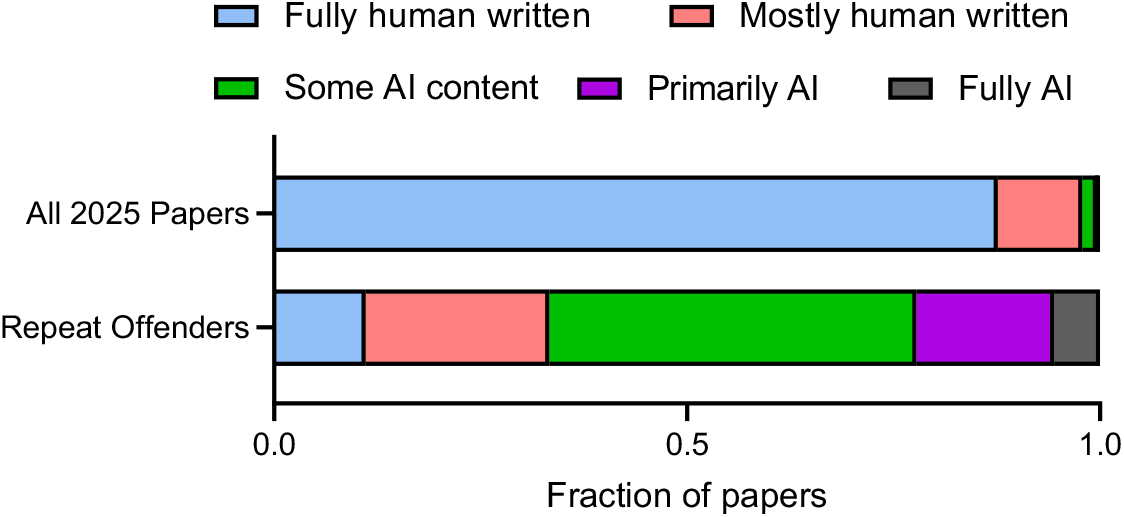
AI-generated content among additional manuscripts published by senior authors of fully AI-generated papers.

Lastly, we tested whether AI-generated writing was more prevalent in some journals than others. Although individual journal policies differ, nearly all require authors to disclose any use of generative AI beyond minor copy editing. In our dataset, papers published in highly selective journals such as *Nature, Science*, and *Cell* contained exceedingly few examples of AI-generated text (Figure 4). In the small number of cases where such text was detected, the AI-positive windows were almost always confined to the final sections of the manuscript, typically the “Limitations of the Study” or concluding remarks within the Discussion section, rather than the main body of the manuscript. In contrast, the two journals with the highest absolute number of AI-positive papers were also the two most prolific publishers in our 2025 dataset, each contributing over 1,000 manuscripts. At this scale, it is plausible that manuscripts containing AI-generated prose may pass through without adequate safeguards, even though peer review in principle should serve as one such check. These patterns are compatible with several behavioral models, including stricter editorial gatekeeping at the most prestigious journals and greater author-driven investment in human editing when preparing manuscripts intended for high-impact venues.

**Figure 4:**
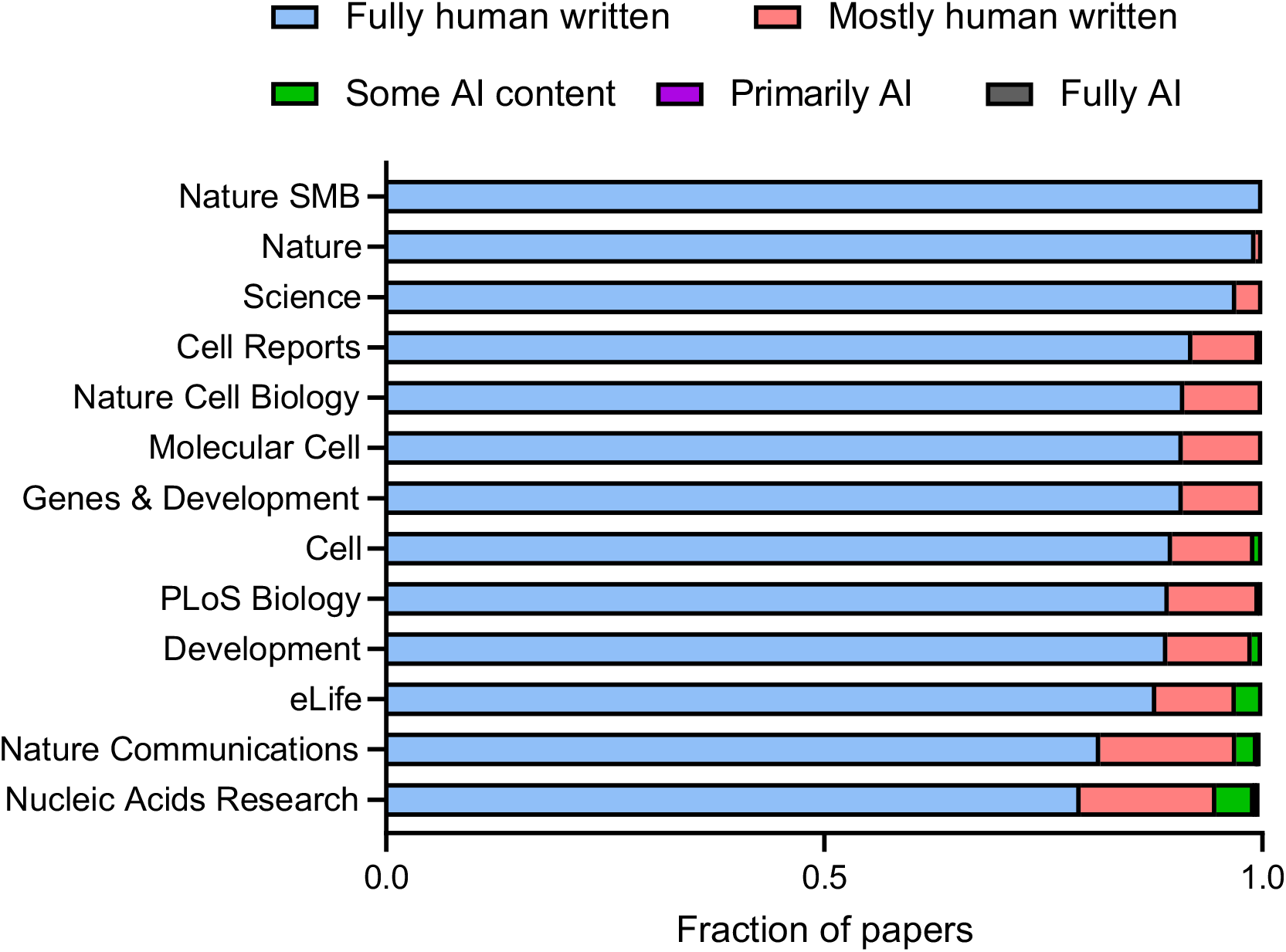
Prevalence of AI-generated text segmented by journal.

## Discussion

Amidst the trillions of dollars of AI investment seeking to reshape the global economy, the architects of these models have set their sights on a loftier and more ambitious goal: the acceleration of science itself. Sam Altman has argued that the main benefits to human welfare from AI will come from faster scientific progress^11^. Demis Hassabis describes DeepMind’s mission as “solving intelligence” in order to advance science and benefit humanity^12^. Dario Amodei goes further, predicting that AI-enabled biology and medicine will “compress the progress that human biologists would have achieved over the next 50–100 years into 5–10 years,” a vision he calls the “compressed 21st century”^13^. These are bold visions. Yet despite their impressive capabilities, current models are not yet capable of independently driving scientific discovery at scale. For now, they largely function as hybrid partners: useful across the day-to-day workflow, but most visibly adopted when researchers turn to the task of writing a scientific manuscript.

Here, we measured the degree to which AI-generated text is already embedded in published manuscripts. Our analysis functions as a natural experiment that reveals how working scientists actually deploy frontier models, and provides a real-world measuring stick of the practical value AI is delivering inside the research pipeline. We observe that significant AI usage only begins in papers published in 2025, but will in all likelihood accelerate in coming years.

The scientific community now faces an unavoidable question: how should we regulate, accommodate, or discourage AI-generated writing in published research? One possible path is a strict prohibition. Journals could adopt a zero-tolerance policy on AI-generated text, motivated by concerns that the scientific literature must remain free of model-generated contamination. If biomedical papers containing AI-generated prose were to be later incorporated into model training corpora, one might reasonably worry that this introduces the same kind of synthetic contamination seen in controlled studies, where models trained heavily on their own outputs reliably degrade in quality^14–16^. A meaningful prohibition, however, would require real enforcement. Retractions would need to be issued when AI-generated content is discovered after publication, and any such system would always lag behind advances in model capability.

A second path is the status quo: journals require disclosure of AI usage, and authors are expected to comply. Our results suggest that this arrangement is untenable in practice. Because AI use carries social stigma, authors rarely disclose it, even when generative text is present. This dynamic fosters a widening gap between stated norms and actual practice, which is corrosive in a moment when public trust in the scientific enterprise is already unusually fragile. If disclosure is to serve as the primary ethical standard, cultural expectations around AI use will need to shift dramatically; at present, they have not.

A third path is the adoption of state-of-the-art AI detectors as part of routine editorial screening. Yet this, too, has structural limitations. Goodhart’s Law states that “when a measure becomes a target, it ceases to be a good measure,” and the rapid iteration of frontier models ensures a moving adversarial boundary. Detectors may flag yesterday’s models reliably while being blind to tomorrow’s. Furthermore, it is not obvious that journals wish to become adversarial enforcement bodies, nor is it clear what the appropriate remedy should be when AI-generated text is detected during peer review. Should the manuscript be rejected outright? Should authors be required to rewrite the entire text? These questions remain unresolved.

Our goal in this study is to provide empirical grounding for a conversation that the scientific community must begin in earnest. Generative AI is sure to continue to shape the way research is conducted, communicated, and evaluated. The challenge ahead is to develop norms that preserve the integrity of the scientific record while enabling the legitimate and potentially transformative uses of these tools. Like all powerful technologies, AI is a double-edged instrument—capable of expanding human creativity and insight, yet also capable of degrading the standards upon which scientific progress depends. The responsibility for navigating this transition falls to us.

## Methods and materials

### Literature corpus construction and text extraction

Biomedical research articles were retrieved from PubMed using the National Center for Biotechnology Information (NCBI) E-utilities application API interface. PubMed identifiers were mapped to PubMed Central identifiers where available, and parsed using custom Python scripts to extract the main narrative text.

### Text preprocessing and cleaning

To focus analyses on author-written scientific prose, we excluded non-narrative and ancillary sections, including Materials and Methods, acknowledgements, references, figure legends, tables, and supplementary material. Inline citations, equations, and formatting artifacts were removed, and paragraph structure was normalized. All text was minimally cleaned to preserve original authorial style while eliminating nonsemantic noise.

### Metadata integration

Article-level metadata, including publication year, journal, author affiliations, and country of origin, were integrated using the OpenAlex database. PubMed and Digital Object Identifier mappings were used to reconcile records across data sources.

### AI-generated text detection

Cleaned article text was analyzed using the Pangram API. Text was segmented into overlapping windows, and each window was assigned an AI-likelihood score. For document-level analyses, we used Pangram’s summary statistic outputs, combining “Mostly human written, with small amount of AI content detected” and “Mostly human written, some AI content detected” into the “Mostly human written” category and renaming “AI content detected, but not fully AI-Generated” as “Some AI content”.

### Statistical analysis and visualization

Statistical summaries were manually plotted, with papers binned by publication year, journal, and geographic origin, and results were visualized using simple bar plots and histograms.

### Use of artificial intelligence tools

Generative artificial intelligence tools were used in during this study. Large language models were used to assist with software development and code refinement. AI tools were also used for editorial assistance during manuscript preparation. All analyses, interpretations, and conclusions were conceived by the authors, and all code and text were reviewed and edited by the authors for accuracy and originality.

## Data availability

The primary data analyzed in this study consist of full-text scientific manuscripts and derived metadata, the redistribution of which is restricted to protect author privacy and to comply with publisher and database usage terms. Processed summary statistics and aggregate results sufficient to reproduce the analyses presented here are included in the manuscript. Additional details regarding data processing and analysis are available from the corresponding author upon reasonable request.

## Acknowledgements

We thank Rachel Matt, James Valcourt, and George Hageman for reading early drafts of this paper. We thank Ryan Briggs for first bringing Pangram to our attention. We thank Katherine Thai, Max Spero, and Bradley Emi for facilitating access to Pangram for large scale analyses.

